# High Throughput Single Cell Proteomic Analysis of Organ Derived Heterogeneous Cell Populations by Nanoflow Dual Trap Single Column Liquid Chromatography

**DOI:** 10.1101/2023.01.06.522908

**Authors:** Simion Kreimer, Aleksandra Binek, Blandine Chazarin, Jae Hyung Cho, Ali Haghani, Alexandre Hutton, Eduardo Marbán, Mitra Mastali, Jesse G Meyer, Thassio Mesquita, Yang Song, Jennifer Van Eyk, Sarah Parker

## Abstract

Identification and proteomic characterization of rare cell types within complex organ derived cell mixtures is best accomplished by label-free quantitative mass spectrometry. High throughput is required to rapidly survey hundreds to thousands of individual cells to adequately represent rare populations. Here we present parallelized nanoflow dual-trap single-column liquid chromatography (nanoDTSC) operating at 15 minutes of total run time per cell with peptides quantified over 11.5 minutes using standard commercial components, thus offering an accessible and efficient LC solution to analyze 96 single-cells per day. At this throughput, nanoDTSC quantified over 1,000 proteins in individual cardiomyocytes and heterogenous populations of single cells from aorta.

**For Table of Contents Only:** 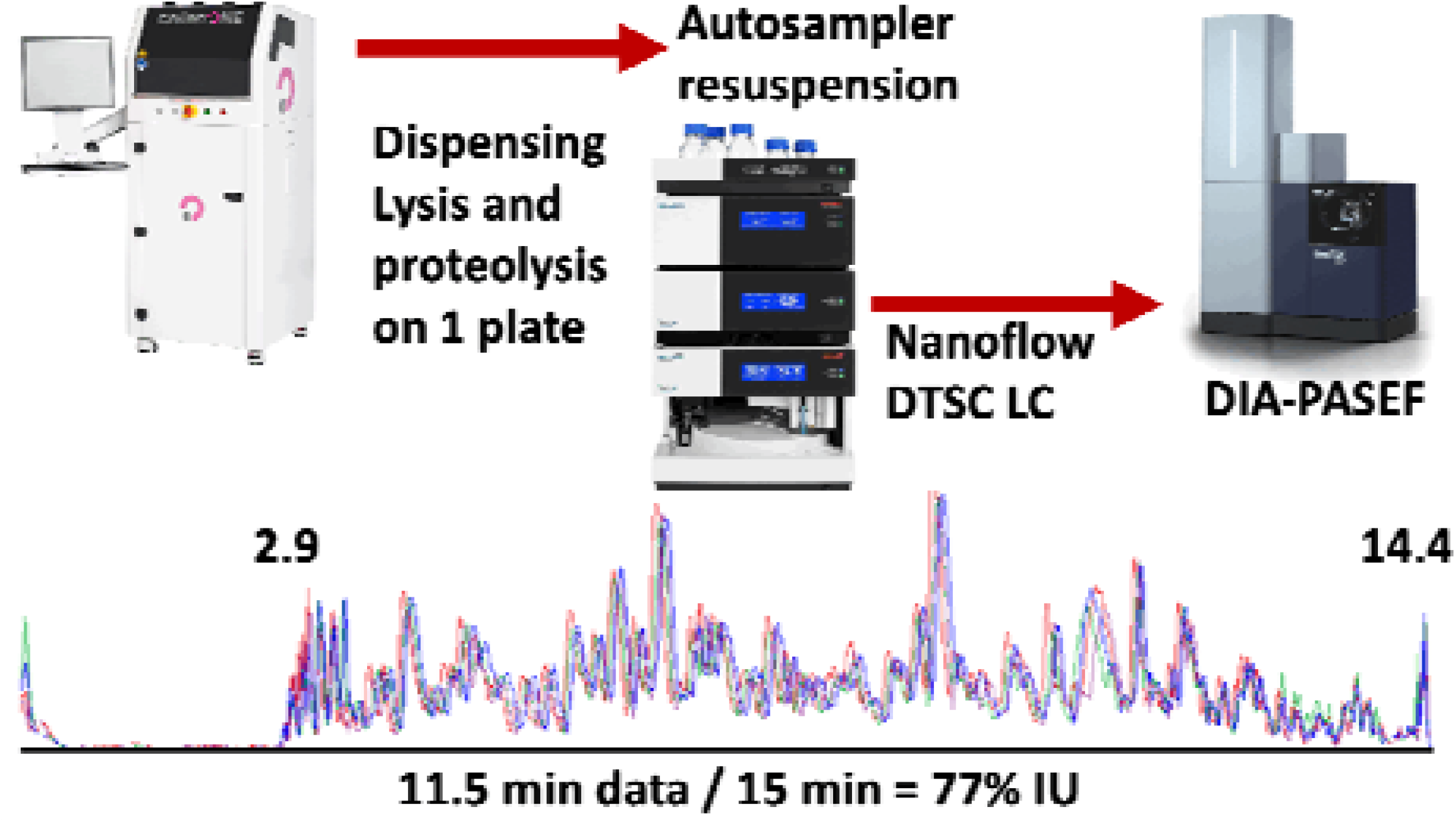

## Introduction

Single cell transcriptomics has revealed unique cell subpopulations within a myriad of tissues^1,2^ but complementary global measurements of the corresponding proteomic phenotypes via LC-MS have been hampered by the limited quantity of protein and comparatively slow analysis. Initially multiplexing by isobaric mass tag (TMT) labelling addressed these challenges by combining analysis of up to 16 samples and increasing sensitivity with a “booster” channel containing a pool of cells (SCoPE-MS).^3,4^ The combined signal from proteins shared between samples in the multiplex set boosts the fragment signals into a reliably identifiable range. Complex samples such as dissociated organs or pathological specimen with substantial heterogeneity in the proteomes of phenotypically distinct cells challenge this strategy because proteins unique to a phenotype are not amplified like shared proteins and thus are less likely to be identified. Reliance on a “booster” introduces bias for the dominant cell type(s) and further reduces the profiling of rare cells.^5,6^ Direct, label-free analysis of individual cells is unbiased in capturing all phenotypes during proteomic analysis of heterogenous tissues and organs.

Label-free analysis of individual cells has been achieved via highly efficient ionization at ultralow flow-rates (*e.g*. 10-30 nL/min)^7-10^ and by improved MS ion transmission and sensitivity.^11^ Analysis at low flow-rates is slowed by the dwell volume of the system and the time spent loading the sample, cleaning, and equilibrating the system resulting in a low amount of data collected relative to total instrument operation time (instrument utilization, IU). Operation of two analytical columns in parallel, where a sample is loaded on one column while a second sample is analyzed elegantly increases IU, but requires a specialized ion source that accommodates two emitters and an LC with two analytical pumps.^9^ Our dual trap single column (DTSC) platform optimized for the nano flowrate (500 nL/min, nanoDTSC) increases the throughput of single cell analysis by parallelizing sensitive high ion accumulation time (166 ms) DIA-PASEF data acquisition on the timsTOF-SCP (Bruker) with resuspension, loading, and desalting of the subsequent sample.^12^ NanoDTSC does not require specialized instrumentation, so it is widely accessible; here an Ultimate 3000 nanoRSLC (Thermo) with the standard Bruker captive spray ionization source were used.

## Experimental Details

### Liquid Chromatography

The dual trap single column configuration (DTSC)^12^ was adapted for single cell analysis (**Fig. S1**) using cartridge trapping columns with a 0.17 μL media bed (EXP2 from Optimize Technologies) packed with 10 μm diameter, 100 Å pore PLRP-S (Agilent) beads, and a PepSep 15 cm x 75 μm analytical column packed with 1.9 μm C18 solid phase (Bruker). The configuration was installed on a Thermo Ultimate 3000 nanoRSLC equipped with one 10-port valve, one 6-port valve, and a nano flow selector. Two complementary methods controlled this configuration. In each method the analytical gradient was delivered through one trapping column to elute the captured peptides unto the analytical column for electrospray ionization into the mass spectrometer while the loading pump flushed the alternate trapping column with organic buffer from the autosampler loop then loaded and desalted the subsequent sample. In the next run, the 10-port valve was switched and the roles of the trapping columns were reversed. The 6-port valve regulated the direction of the loading pump flow through each trapping column to backward flush and remove precipitates at the trapping column head and forward load the sample to prevent any precipitate from reaching the analytical column. (**Fig. 1A**).

**Figure 1.**
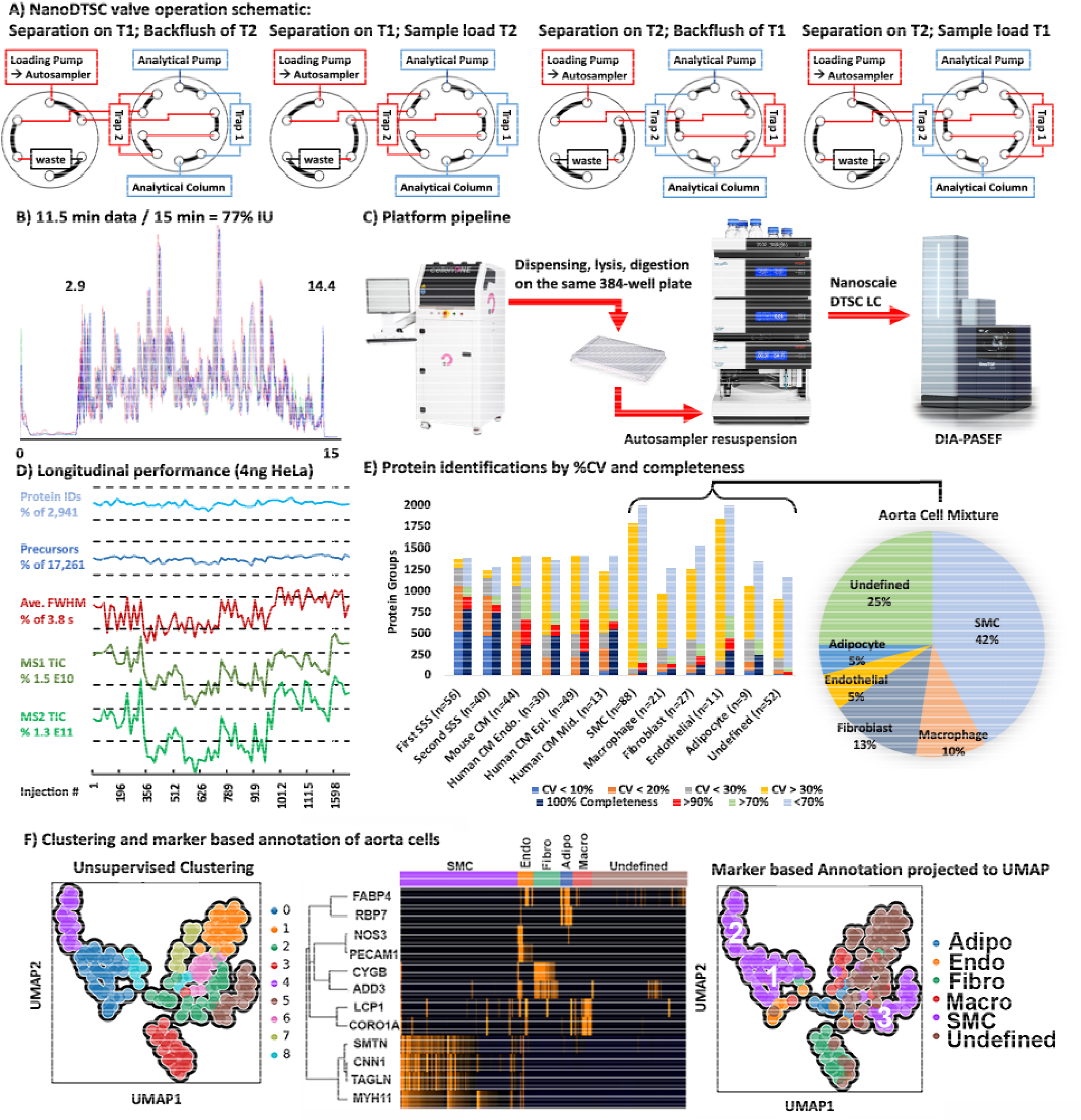
A – Valve schematic of the two methods used in nanoDTSC blue represents the analytical separation path and red represents the path of loading pump flow used for cleaning, equilibration, and sample loading. B – Overlay of 4-SSS runs, peptides elution 2.9-14.4 minutes (77% instrument utilization). C – Workflow overview: Individual cells are dispensed by CellenOne into 384-well plate, digested, resuspended by the autosampler, and analyzed by DIA-PASEF. D – Consistent system performance over 1,700 injections monitored by HeLa QC, dashed lines represent 10% deviation from average. E –Protein quatitative reproducibility with color coded %CV ranges (blue <10%, orange <20%, grey <30%, yellow >30%) and color coded data completeness (dark blue detected in every sample, red >90% of samples, light green >70% of samples, grey <70% of samples) for each sample type analyzed. The distribution of annotated cells from aorta is represented as a pie chart with smooth muscle cells comprising the largest group. F – Unsupervised UMAP clustering of aorta cells by entire proteome (left), clustering by cell-type markers (middle), and projection of cell type annotation unto the UMAP plot (right).

The analytical separation used 0.1% formic acid in water for mobile phase A and 0.1% formic acid in acetonitrile for mobile phase B and the gradient was delivered as follows (**Fig. S1**): start at 9% B at 500 nL/min; linear increase to 22% B over 8 minutes; linear increase to 37% B over 4.7 minutes; increase to 1000 nL/min flowrate and 98% B over 0.2 minutes; hold at 98% B for 1 minute; drop to 9%B at 1,000 nL/min over 0.1 minutes; hold at 9% B at 1,000 nL/min for 0.9 min; return to 500 nL/min flowrate in 0.1 min (15 min total). The loading pump delivered 0.1% formic acid in water at 20 μL/min for the first 6 min of the run during trapping column cleaning, and dropped to 10 μL/min from 6.5 to 15 min to load and desalt the subsequent sample. The valves and trapping columns were kept at 55° C in the Ultimate 3000 column oven compartment and the analytical column was kept at 60° C using the Bruker “Toaster” oven.

The autosampler was programmed to immediately initiate data acquisition, then collect acetonitrile with 0.1% formic acid into the 20 μL sample loop and inject it through the trapping column using the loading pump flow. After repeating the acetonitrile flush a second time, the loop and needle assembly were rinsed with 25 μL of 0.1% formic acid in water and the autosampler aspirated 20 μL of 2% acetonitrile in water with 0.1% formic acid. This solvent was dispensed and aspirated into the sample well three times to resuspend the sample. The sample was injected and the needle assembly was rinsed (**Fig. S1**).

### Mass Spectrometry

The analytical column was directly connected to a 20 μm ZDV emitter (Bruker) installed in the Bruker captive source. The capillary voltage was set to 1700 V with the dry gas set at 3.0 L/min and 200° C. Data were acquired by DIA-PASEF with the ion accumulation and trapped ion mobility ramp set to 166 ms. DIA scans were acquired with 90 m/z windows spanning 300-1200 m/z and 0.6-1.43 1/K0. One full MS1 scan followed by 4 trapped ion mobility ramps to fragment all ions within this region resulted in a 0.86 s cycle time. The DIA windows are presented in **Fig. S2**.

### Sample Preparation

Individual cells were dispensed into separate wells on a 384-well low binding PCR plate (Biorad HSP3801) containing 600 nL of lysis buffer using a CellenONE (Cellenion) as previously reported by the Mechtler lab.^13^ Samples were not treated with alkylating or reducing agents. After lysis by freeze-thaw the samples were subjected to trypsinization with 50 ng/μL trypsin, followed by acidification using a protocol adapted from one-pot sample preparation method.^3^

### Data Analysis

Data were analyzed with DIA-NN 1.8.1^14,15^ using sample specific libraries for each sample type. Each search was conducted with second-pass and match-between-runs (MBR) enabled. The mass error tolerances were set at 15 ppm for both fragment and intact masses.

An initial library was generated for mouse cardiomyocytes and the mouse heart tissue digest using gas phase fractionated (GPF) DDA-PASEF in which the selection of precursors for MS2 was limited to 300 m/z ranges spanning 300-1200 m/z (*i.e*. 300-600 m/z, 600-900 m/z, *etc*.). Aside from the data acquisition scheme the library generating methods were identical to the methods used to analyze individual cells by DIA. Mouse cardiomyocytes and heart tissue lysates were analyzed over 5 replicates at each m/z range and 5 replicates for the full 300-1200 m/z range (50 runs total) and searched against the mouse protein database and compiled into an EasyPQP library using FragPipe (18.0). This initial GPF-DDA library was used to analyze the DIA data from mouse cardiomyocytes and heart tissue replicates. The 10 cell and 10 heart tissue DIA runs with the highest identifications were analyzed using the library-free search in DIA-NN against the mouse protein database. The library-free identifications at < 1% FDR were combined with the DDA based identifications to generate the final merged library used for analysis of mouse CMs and heart tissue samples. It should be noted that combining libraries also combines the false positive entries, so the real FDR for this combined cardiomyocyte library is 1-2%. The two libraries contained complimentary populations of precursors with only a 43% overlap, and searches with the merged library on average identified 16% more precursors than individual libraries (**Fig. S3**).

The strategy was repeated for analysis of mouse aorta extracted cells. The macrophage cells were not well represented in the GPF DDA-PASEF library so data from PXD024658^6^ was reanalyzed so that spectra could be used to supplement the library. Again, the ten cells with the highest identifications from each putative type were analyzed using the library-free strategy to add DIA-based identifications. The library compiled from the three sources had an FDR of 1-3%.

No GPF DDA-PASEF library was generated for the human cardiomyocytes and the entire cohort was analyzed only using the library-free feature of DIA-NN. The identification false discovery rate in cell analysis was filtered to 1% by DIA-NN using the target-decoy strategy.

Protein level quantitation was used to cluster individual cells using the Python Single-Cell Analysis package (SCANPY)^16^, which models the data as a connected graph. The graph is constructed using Uniform Manifold Approximation and Projection (UMAP).^17^ UMAP identifies the *k* (set to 12) nearest neighbors for each cell, uses those neighbors to compute the cell’s local connectivity, then combines the local connectivity to form a graph representation of the data. Leiden clustering^18,19^ was applied to the resulting graph to identify neighborhoods, or clusters within the graph. For this paper, we used 12 neighbors to compute local connectivity, did not exclude cells/proteins, and used only the presence of proteins to determine their relevance. We did not consider the absence of proteins in determination of relevance.

### Data Availability

The presented data is shared on MassIVE : MSV000090903,<u>PXD038828</u>,https://doi.org/doi:10.25345/C5Z60C65X, password: dualtrapscp.

## Results and Discussion

The developed method collected 11.5 minutes of data (span of peptide elution) during 15 minutes of total run-time with negligible delay (2 s) between runs to achieve 77% IU (**Fig. 1A**). Sample loss was minimized by dispensing,^2^ lysing, and digesting cells with a modified one-pot protocol^20^ on the same 384-well plate in 600 nL of solvent. The generated peptides dried rapidly and were resuspended and injected by the LC autosampler (**Fig. 1B**). The 5-10 minutes required for thorough sample resuspension and loading were recovered through parallelization which increased throughput by at least 25% compared to conventional trap and elute with the same components. In this implementation a flowrate of 500 nL/min was used to reduce dwell time (the 2.9 min required for trapped peptides to elute from the analytical column). Operation at a lower flowrate should be more sensitive but the efficient ion transmission of the mass spectrometer and the increased ion accumulation time (166 ms) still achieved an adequate depth of profiling. The implemented method thoroughly cleaned the 0.17 μL trapping columns with two 20 μL (∼100 column volume equivalents) loop injections of acetonitrile. Hence carryover was negligible as determined by identification of 0 proteins in blank injections directly following cell analysis.

Method reproducibility and robustness were evaluated using 3 ng of bulk digested mouse heart tissue as the system suitability sample (SSS). Analysis of 56 technical replicates quantified on average 1,145 mouse cardiac proteins with 1,063 at high precision (coefficient of variance, CV < 20%). After almost 1,700 injections using the same trapping and analytical columns a second batch of 40 SSS injections was analyzed and an average of 1,025 proteins were quantified, with 925 proteins under 20% CV demonstrating a minor loss in performance after 3 weeks of continuous operation. Longitudinal system performance was also monitored by analysis of 4 ng of Pierce HeLa standard once on each trap every 50 single cell runs. On average 17,290 precursor ions and 2,942 proteins were identified without significant deviation (< 10% deviation from average) through the course of analysis. Peak widths and ion current MS1 and MS2 intensities also remained stable (< 20% deviation from average, **Fig. 1C**). A relatively homogenous population of heart extracted cardiomyocytes and a heterogenous mixture of cells from mouse aorta were used test the utility of this platform. Analysis of 44 individual wild-type mouse cardiomyocytes found 96.9% proteome overlap with the SSS and similar numbers of identified precursors and proteins. The reproducibility and data completeness were lower between individual mouse cardiomyocytes than the SSS replicates due to biological heterogeneity. Human cardiomyocytes isolated from the endomyocardium (n = 30), midmyocardium (n = 13), and epimyocardium (n = 49) of the left ventricle of one donor heart had similar protein identification numbers and data completeness but even higher quantitative heterogeneity (**Fig. 1D**). This variance could be attributed to the wide cell size distribution (CellenOne diameter: 47 - 114 μm), as it exceeds the technical imprecision baseline established from SSS and mouse cardiomyocyte analysis. The rank vs. log intensity plot of heart tissue and cardiomyocytes shows highly similar protein identifications and a distribution spanning roughly 5 orders of magnitude with high data completeness that drops off for lower abundance proteins (**Fig. S4A**). Dimension reduction of protein quantities from human cardiomyocytes with Uniform manifold approximation and projection (UMAP) found that the number of identified proteins was the biggest driver of clustering. Separate clustering of the high identification (>1100 proteins) and low identification groups (<1100) still did not identify clear subpopulations or differences in source tissue layers (**Fig. S5**). The ability to identify cardiomyocyte subpopulations by UMAP is hindered by the low cell number (92) and the quantitative variance resulting from the wide cell size distribution.

Conventional normalization strategies such as adjustment based on median or total protein that are typically used to reduce variance due to differences in bulk protein content may not be appropriate for cells spanning a range of sizes, lifecycle stages, and sub-types. We attempted a ratio-based analysis where each protein is divided by every other protein within each cell (limited to proteins quantified in all cells, 255 for this set) and the CV of each ratio across the cell population is used to infer correlations in protein expression. In human cardiomyocytes, 4,010 ratios from 154 proteins had a CV under 20%. These proteins were from mitochondrial complexes which is not surprising since their relative abundance is independent of the number of mitochondria within a cell and tightly regulated. Proteins from the sarcomeric complex appeared in the 11,000 ratios from 204 proteins within the 20-30% CV range (**Fig. S6**). In agreement with known sarcomeric protein complex composition,^21^ the average proportion of Myosin light chain (MYL3) to heavy chain (MYH7) was 0.5 (25% CV). Additionally, myosin light chain 2 (MYL2) to myosin light chain 3 (MYL3) was a stable 1:1 ratio with (CV 30%) and the proportion of tropomyosin 1 (TPM 1) to tropomyosin 2 (TPM2) was also a stable average of 8.4 (26% CV) (**Fig. S4B**). Ratio-based analysis of mouse smooth muscle cells (SMC) extracted from aorta also found high regulation of mitochondrial protein ratios and reproducible ratios of tropomyosin 1 to tropomyosin 2 (1.3, CV 16%). Interestingly, SMC heavy (MYH11) to regulatory light chain (MYL9) ratio was on average 7.5 with substantial heterogeneity between SMC subtypes (CV 31%, subtype annotation described below) (**Fig. S7**). This analysis reflects fundamental biology and circumvents the challenge of individual protein intensity heterogeneity due to cell size distribution, but it should be noted that ratios combine quantitative imprecision of each protein and are inflated by outlier cells. Furthermore, the coefficient of variance is not an adequate metric in cell populations where the protein quantities are regulated only in some of the cells or are expressed in two or more ratios. More sophisticated statistical methods that identify outliers and multiple distributions are needed in future work. By identifying stable ratios within this healthy population of cardiomyocytes it is possible to identify disease or novel phenotypes based on disruption of these ratios. Detection of cardiomyocytes with unusual ratios of mitochondrial or sarcomeric proteins in a biopsy would be a more sensitive diagnostic than bulk tissue analysis in which these changes would be masked by healthy cells.

The aorta contains a heterogeneous mixture of smooth muscle cells (SMC), adipocytes, endothelial cells, fibroblasts, macrophages, and other immune cells. Cells from dissociated mouse aorta were analyzed by nanoDTSC and putatively annotated and clustered based on the presence of known markers. Dimensionality of quantitative single cell protein data was reduced by UMAP and clustered with the Leiden algorithm. Putative marker-based annotations were projected onto the points in the dimension reduced space. For SMCs the annotation mapped unto four unsupervised clusters (#0, 4, 5, and 8) (**Fig. 1F**). Cluster 8 was close to 0 and only a few cells in size, thus we grouped SMCs into 3 proteome-based clusters. Notably, transgelin (SM22alpha) showed the most profound difference in abundance in cluster 5 (SMC3) which is consistent with modified, proliferative SMC phenotype that is biologically significant in disease states (**Fig. S7**).^22^ Fibroblast markers mostly mapped unto a single cluster (#3), which also contained un-annotated cells, suggesting that the proteins used for annotation (CYGB and ADD3) do not represent all fibroblasts and these unannotated cells were assigned as fibroblasts due to overall proteome similarity. Adipocytes, endothelial, and macrophage markers did not map neatly unto clusters, although some patterns could be discerned. Overall, unsupervised clustering is a viable initial strategy for definition of putative cell types in complex mixtures without *a priori* knowledge, and could be improved with higher total cell numbers that include more instances of the less abundant cell-types.

## Conclusion

NanoDTSC reproducibly quantified the proteomes of individual cardiomyocytes to a depth of over 1,000 protein groups in 15 minutes of total run time, with meaningful data acquired for 77% of the time. Longitudinal quality control showed that the system remained robust over 1,700 samples without replacement of components. The platform proved suitable for analysis of heterogeneous cell mixtures extracted from an organ and the combination of cell marker-based annotation and unsupervised clustering was able to define putative cell types. The proposed ratio-based analysis circumvented the challenge of wide cell-size distributions and confirmed the expected stability of mitochondrial and sarcomeric protein complexes. NanoDTSC neatly integrated with automated isolation and tryptic digestion of cells by resuspending the generated peptides from the same 384-well plate unto which the living cells were dispensed thus minimizing sample loss. While 15 minute/cell is adequate throughput for most applications it can be increased further with a lower volume analytical column (**Fig. S8)** and non-isobaric labeled multiplexing (plexDIA).^23^

## Supporting Information

Gradient and autosampler and valve programs (**Fig. S1**); DIA isolation windows (**Fig. S2**); Comparison of libraries for DIA analysis (**Fig. S3**); Protein quantiation in heart tissue and cardiomyocytes (**Fig. S4**); UMAP clustering of cardiomyocytes (**Fig. S5**); Stable protein ratios in cardiomyocytes (**Fig. S6**); Structural protein signatures of SMC subtypes (**Fig. S7**); Initial benchmarks for 10 min/cell nanoDTSC method (**Fig. S8**).

## Acknowledgements

JGM and Alexandre H were supported by the NIGMS (R35GM142502). SP was supported by OWHR/NHLBI R01HL165471.

