## Supplement Information for "High Throughput Single Cell Proteomic Analysis of Organ Derived Heterogeneous Cell Populations by Nanoflow Dual Trap Single Column Liquid Chromatography"

**Table of Contents**

Figure S1. Analytical gradient, autosampler control program, and valve operation	S2

Figure S2. DIA isolation windows	S3

Figure S3. Comparison and combination of libraries for DIA analysis	S4

Figure S4. Protein quantiation in heart tissue and cardiomyocytes	S5

Figure S5. UMAP clustering of cardiomyocytes	S6

Figure S6. Protein ratios below 20% and 30% CV in cardiomyocytes	S7

Figure S7. Distribution of ratios of sarcomeric proteins in SMC	S8

Figure S8. Initial Benchmarks for 10 min/cell nanoDTSC Method	S9

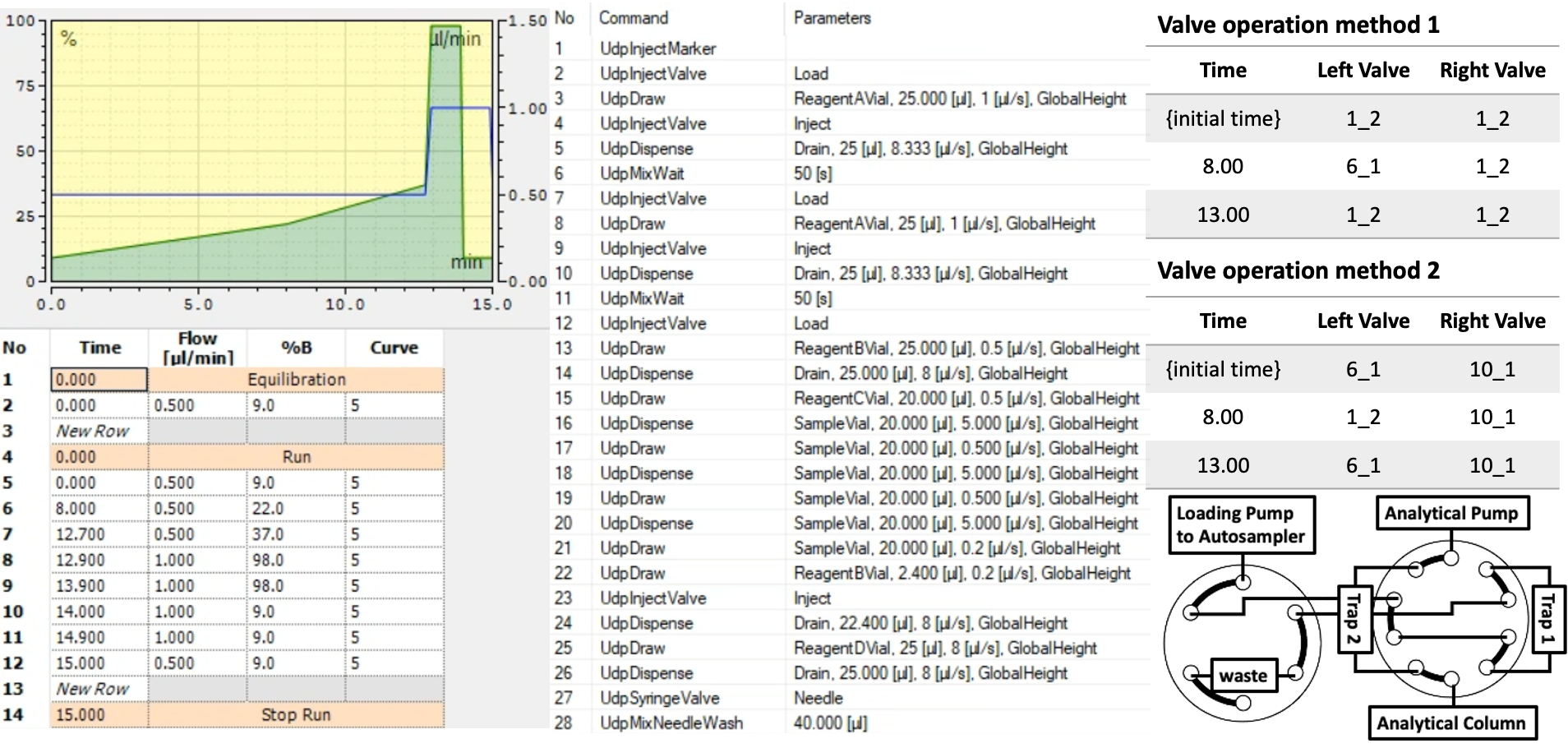

**Figure S1.** Analytical gradient (left), autosampler program (middle), valve operation (right).

| #MS Type | Cycle Id | Start IM [1/K0] | End IM [1/K0] | Start Mass [m/z] | End Mass [m/z] | CE [eV] |
| --- | --- | --- | --- | --- | --- | --- |
| MS1 | 0 | - | - | - | - | - |
| PASEF | 1 | 1.034 | 1.286 | 930 | 1020 | - |
| PASEF | 1 | 0.786 | 0.998 | 570 | 660 | - |
| PASEF | 1 | 0.6 | 0.782 | 300 | 390 | - |
| PASEF | 2 | 1.158 | 1.43 | 1110 | 1200 | - |
| PASEF | 2 | 0.91 | 1.142 | 750 | 840 | - |
| PASEF | 2 | 0.662 | 0.854 | 390 | 480 | - |
| PASEF | 3 | 0.972 | 1.214 | 840 | 930 | - |
| PASEF | 3 | 0.724 | 0.926 | 480 | 570 | - |
| PASEF | 4 | 1.096 | 1.358 | 1020 | 1110 | - |
| PASEF | 4 | 0.848 | 1.07 | 660 | 750 | - |

**Figure S2.** DIA isolation window scheme for DIA-PASEF using 166 ms ion accumulation time and 90 m/z isolation windows.

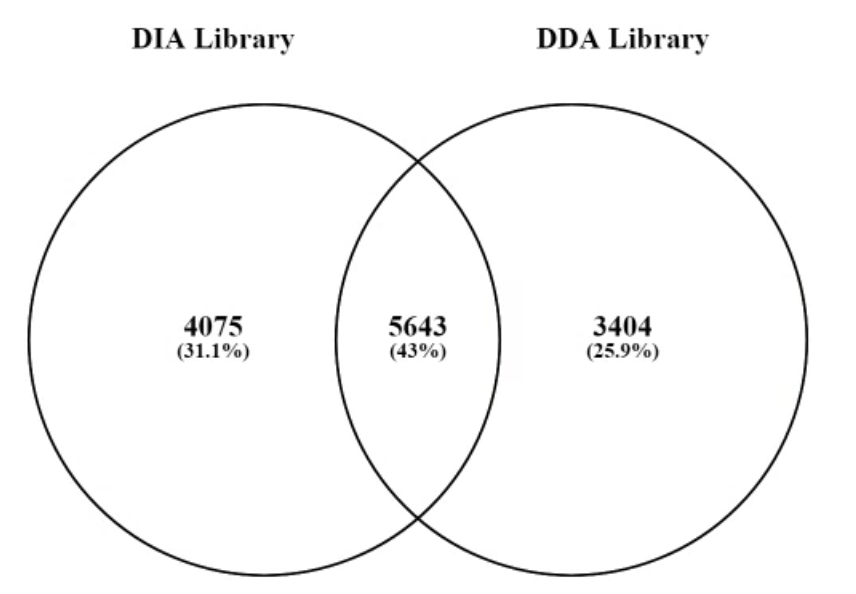

**Cardiomyocytes	3ng Heart Tissue**

Precursor Identifications with Different Libraries

|  | Merged | DIA | DDA |
| --- | --- | --- | --- |
| CM | 8475 | 7547 | 6941 |
| CM | 7860 | 7342 | 6665 |
| CM | 8521 | 7583 | 7020 |
| CM | 9331 | 7922 | 7344 |
| CM | 6433 | 5707 | 5761 |
| CM | 9315 | 7860 | 7221 |
| CM | 9192 | 7914 | 7259 |
| CM | 9327 | 7951 | 7289 |
| CM | 6896 | 6132 | 5724 |
| CM | 9582 | 7935 | 7226 |
| CM | 8064 | 7396 | 6647 |
| CM | 9410 | 7929 | 7289 |
| CM | 9544 | 7867 | 7236 |
| CM | 9128 | 7873 | 7201 |
| CM | 8348 | 7526 | 6882 |
| CM | 9457 | 7928 | 7264 |
| CM | 8775 | 7756 | 7108 |
| CM | 8692 | 7722 | 7015 |
| CM | 8915 | 7809 | 7054 |
| CM | 7312 | 6355 | 5923 |
| CM | 7775 | 7284 | 6719 |
| CM | 8404 | 7574 | 6922 |
| CM | 7044 | 6357 | 5938 |
| CM | 7741 | 7366 | 6644 |
| CM | 8965 | 7796 | 7048 |
| CM | 8485 | 7670 | 6925 |
| CM | 7988 | 7377 | 6707 |
| CM | 7258 | 6958 | 6281 |
| CM | 8872 | 7714 | 6903 |
| CM | 8632 | 7744 | 6992 |
| CM | 8030 | 7237 | 6694 |
| CM | 8460 | 7710 | 6949 |
| CM | 9451 | 7954 | 7187 |
| CM | 6818 | 6554 | 6195 |
| CM | 8248 | 7617 | 6881 |
| CM | 6719 | 6123 | 5964 |
| CM | 7523 | 6943 | 6258 |
| CM | 8888 | 7771 | 6945 |
| CM | 7559 | 6776 | 6156 |
| CM | 7320 | 6622 | 5929 |
| 3ng Heart Tissue | 8604 | 6763 | 7620 |
| 3ng Heart Tissue | 8521 | 6725 | 7534 |
| 3ng Heart Tissue | 8277 | 6560 | 7433 |
| 3ng Heart Tissue | 8468 | 6675 | 7707 |
| 3ng Heart Tissue | 8613 | 6725 | 7638 |
| 3ng Heart Tissue | 8385 | 6603 | 7446 |
| 3ng Heart Tissue | 8370 | 6514 | 7406 |
| 3ng Heart Tissue | 8273 | 6495 | 7400 |
| 3ng Heart Tissue | 7242 | 5542 | 6227 |
| 3ng Heart Tissue | 7669 | 6022 | 6747 |
| 3ng Heart Tissue | 8517 | 6750 | 7573 |
| 3ng Heart Tissue | 7880 | 6073 | 7084 |
| 3ng Heart Tissue | 7649 | 6129 | 6969 |
| 3ng Heart Tissue | 8254 | 6347 | 7378 |
| 3ng Heart Tissue | 8448 | 6701 | 7400 |
| 3ng Heart Tissue | 7762 | 6048 | 6993 |
| 3ng Heart Tissue | 7832 | 6241 | 6957 |
| 3ng Heart Tissue | 8313 | 6439 | 7278 |
| 3ng Heart Tissue | 7538 | 6018 | 6830 |
| 3ng Heart Tissue | 7610 | 6011 | 6934 |
| 3ng Heart Tissue | 8087 | 6378 | 7248 |
| 3ng Heart Tissue | 7490 | 6008 | 6920 |
| 3ng Heart Tissue | 7972 | 6196 | 7052 |
| 3ng Heart Tissue | 7614 | 5982 | 6932 |
| 3ng Heart Tissue | 8157 | 6357 | 7301 |
| 3ng Heart Tissue | 7428 | 5748 | 6490 |
| 3ng Heart Tissue | 7576 | 6056 | 6889 |
| 3ng Heart Tissue | 8022 | 6368 | 7280 |
| 3ng Heart Tissue | 6843 | 5531 | 6453 |
| 3ng Heart Tissue | 7617 | 6101 | 6796 |
| 3ng Heart Tissue | 7065 | 5618 | 6486 |
| 3ng Heart Tissue | 7461 | 5909 | 6751 |
| 3ng Heart Tissue | 7803 | 6304 | 7200 |
| 3ng Heart Tissue | 8261 | 6475 | 7376 |
| 3ng Heart Tissue | 8237 | 6571 | 7468 |
| 3ng Heart Tissue | 7687 | 6262 | 7239 |
| 3ng Heart Tissue | 8142 | 6604 | 7345 |
| 3ng Heart Tissue | 8008 | 6451 | 7401 |
| 3ng Heart Tissue | 8114 | 6520 | 7450 |
| 3ng Heart Tissue | 7884 | 6315 | 7263 |
| 3ng Heart Tissue | 8472 | 6625 | 7541 |
| 3ng Heart Tissue | 7955 | 6241 | 7213 |
| 3ng Heart Tissue | 8005 | 6162 | 7187 |
| 3ng Heart Tissue | 7662 | 6045 | 7044 |
| 3ng Heart Tissue | 7000 | 5768 | 6359 |
| 3ng Heart Tissue | 7839 | 6224 | 7101 |
| 3ng Heart Tissue | 8255 | 6527 | 7339 |
| 3ng Heart Tissue | 7713 | 6181 | 6982 |
| 3ng Heart Tissue | 7939 | 6270 | 7143 |
| 3ng Heart Tissue | 7677 | 6090 | 7009 |
| 3ng Heart Tissue | 7328 | 5765 | 6683 |
| 3ng Heart Tissue | 7034 | 5578 | 6465 |
| 3ng Heart Tissue | 7809 | 6051 | 6999 |
| 3ng Heart Tissue | 6937 | 5364 | 6124 |
| 3ng Heart Tissue | 6864 | 5277 | 5884 |

**Figure S3.** Overlap of libraries generated by gas phase fractionated DDA-PASEF and library-free analysis of the DIA data. The two approaches identify a roughly equal but complementary population of precursors (Venn Diagram). Using the merged library identifies on average 16% more precursors than using either library separately.

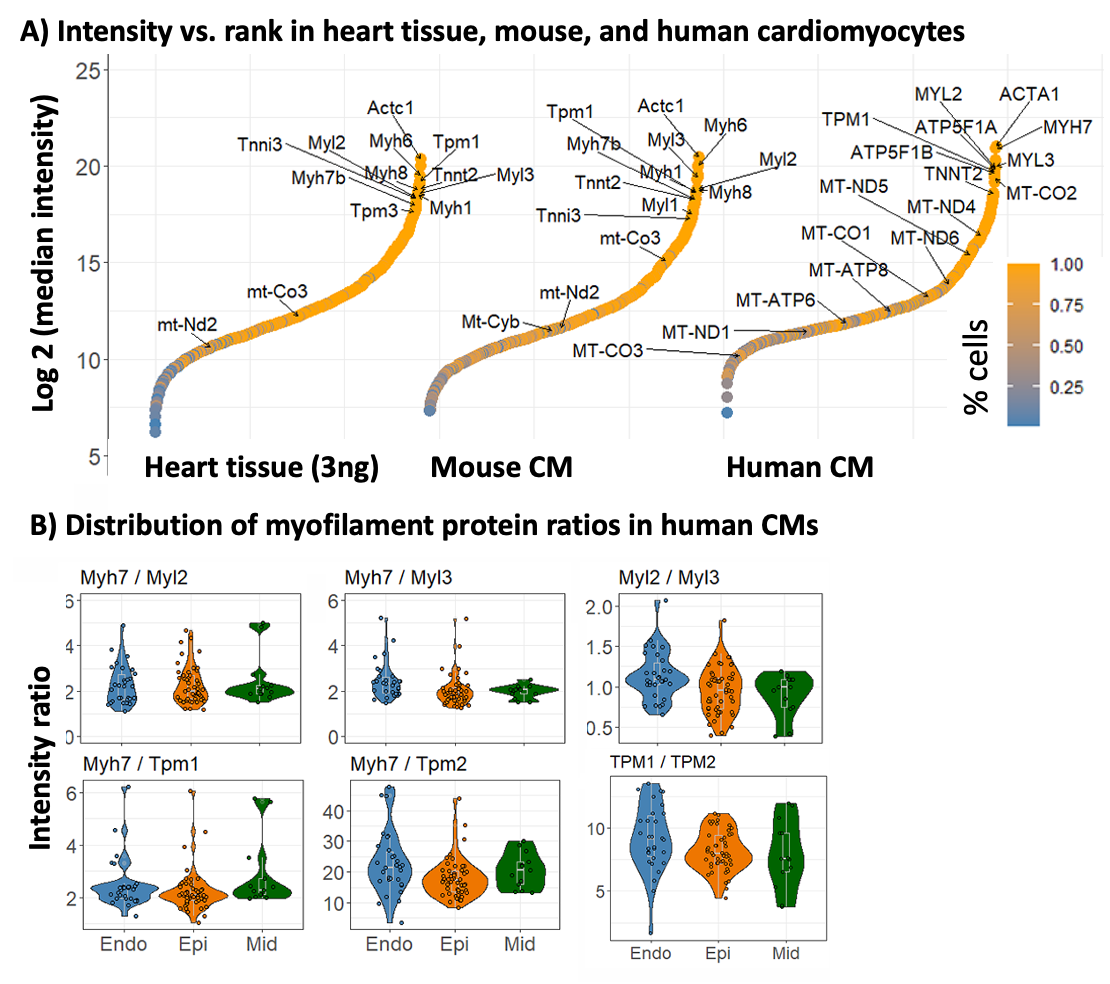

**Figure S4.** A – Log intensity (y-axis) vs rank (left – least abundant left to right – most abundant) curves, data completeness is indicated by color with orange proteins detected in all samples. B – Ratios of sarcomeric proteins in human cardiomyocytes.

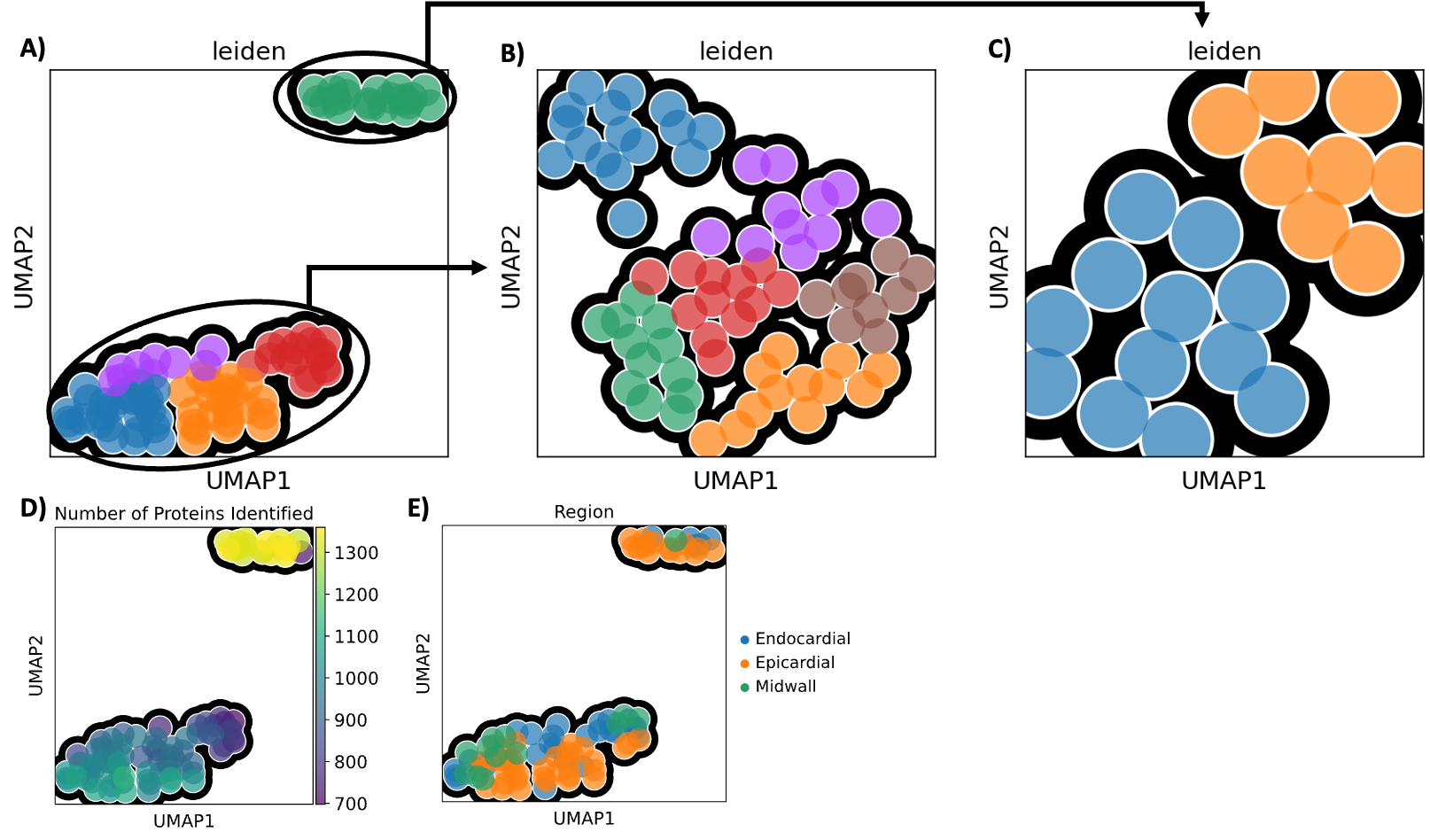

**Figure S5.** A – UMAP plots of all human cardiomyocytes together. B – Cardiomyocytes with less than 1,100 protein identifications. C – Cardiomyocytes with more than 1,100 proteins. D – Projections of protein identifications. E – Projection of region of cell extraction.

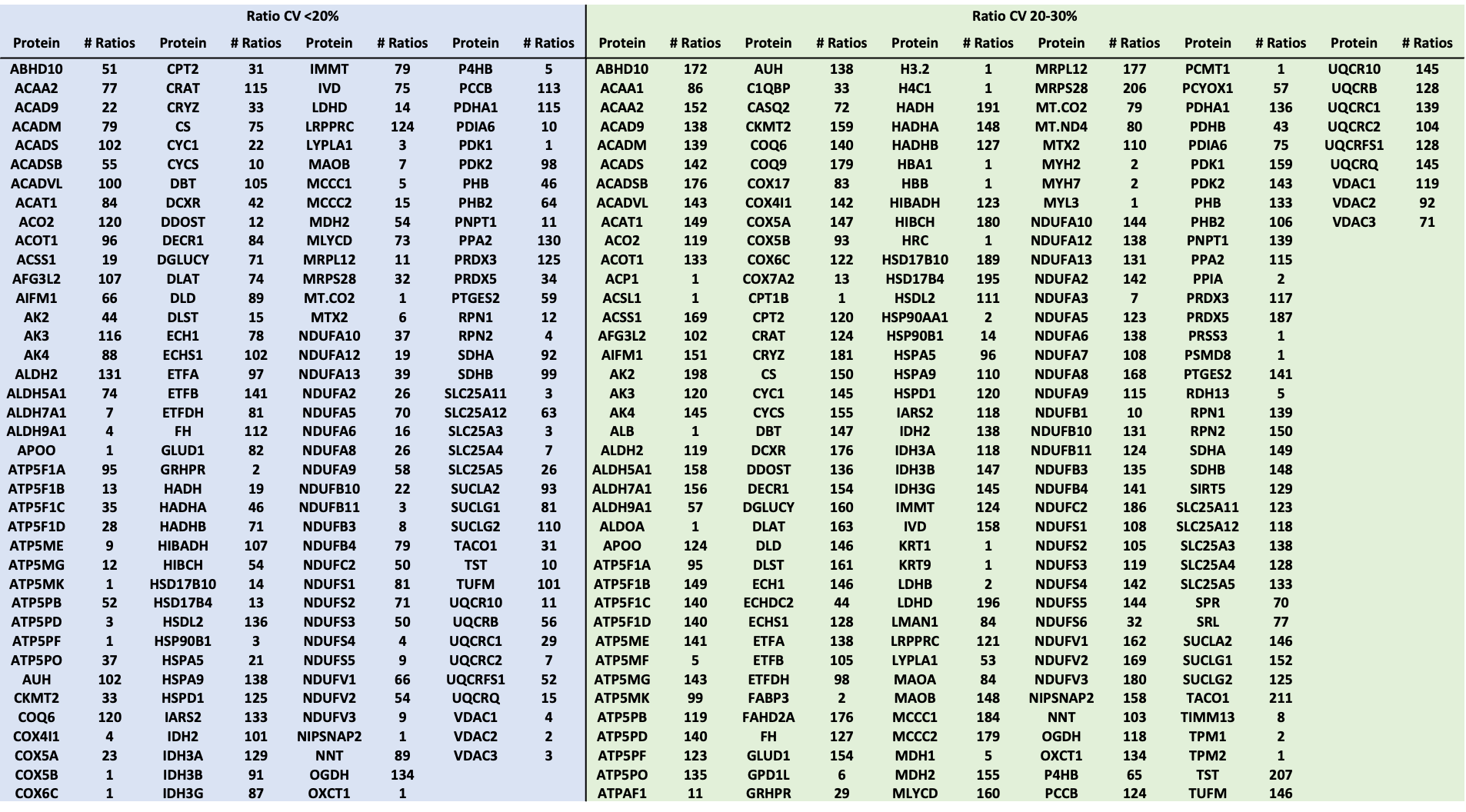

**Figure S6.** Table of proteins involved in stable (CV <20% or 20-30%) ratios across the human CM population and the ratio count.

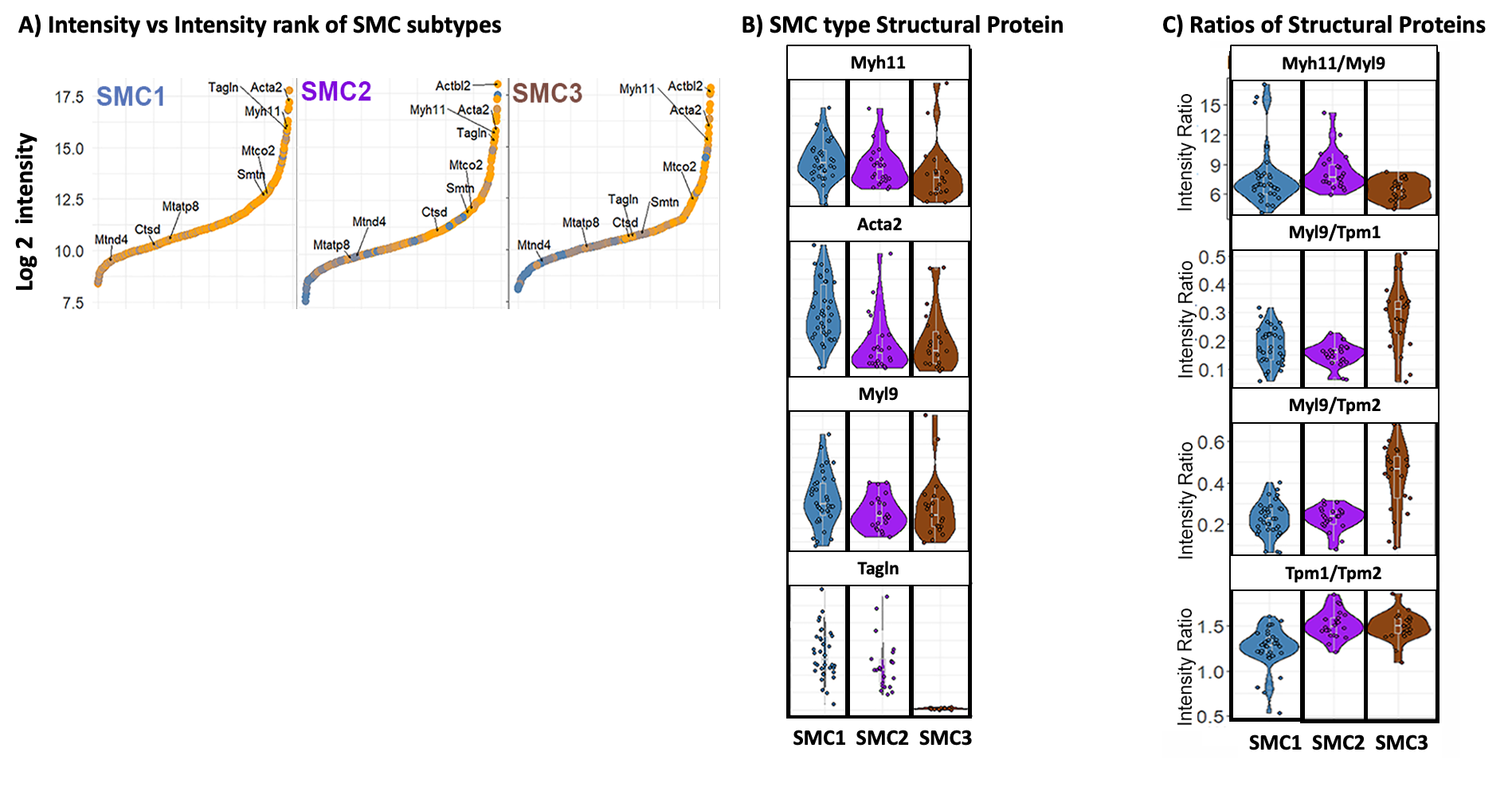

**Figure S7.** A – Distribution of proteins in the three SMC subtypes. B – Abundances of structural proteins distinguishing the three SMC subtypes. C – Stable ratios of myofilament proteins in the three smooth muscle cell subtypes also distinguish subtypes.

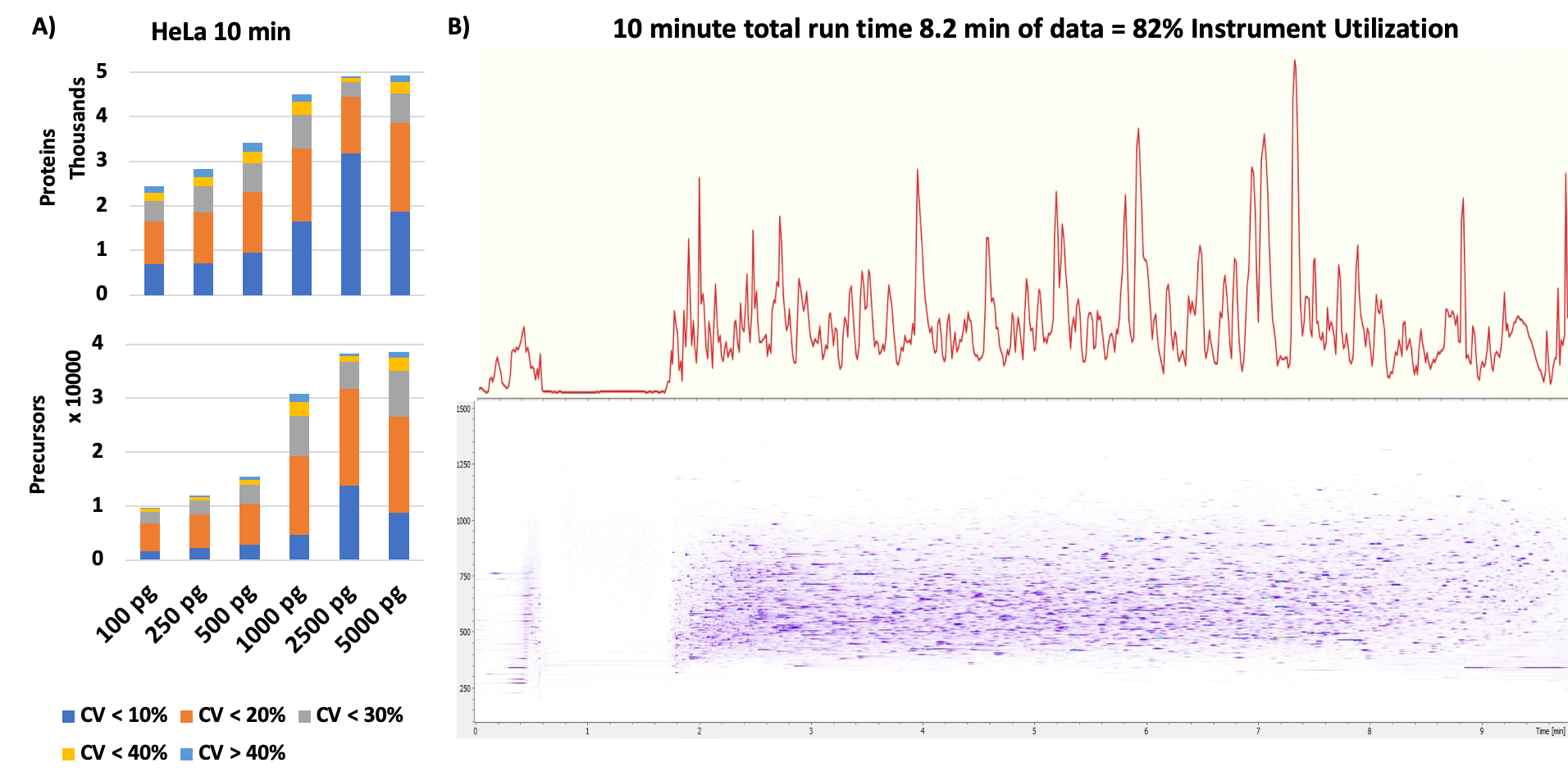

**Figure S8.** A – Preliminary benchmarks with varying HeLa loads using a Aurora Rapid 75 (IonOpticks) column 50 x 0.75 mm, C18 1.7 µm beads and 10 minute total run time at 600 nL/min flowrate. B – Chromatogram and heatmap of same run.
